# Reverse genetics system for emerging tick-borne orthonairovirus highlights dispensable N-terminal region of Gn associated with replication in tick vectors

**DOI:** 10.64898/2026.09.28.755240

**Authors:** Yume Mimura, Takahiro Hiono, Hiromu Arakawa, Shiori Go, Asako Shigeno, Mebuki Ito, Eri Fujii, Keita Mizuma, Ryo Nakao, Yasuko Orba, Hiroyuki Kaji, Keita Matsuno

## Abstract

A growing number of emerging and re-emerging tick-borne orthonairoviruses belonging to the Sulina and Tamdy genogroups have recently been identified in association with febrile diseases in East Asia. Yezo virus (YEZV) is one such viruses and is genetically distinct from well-characterized Crimean-Congo hemorrhagic fever virus. Orthonairovirus glycoproteins (Gn and Gc) expressed from a single glycoprotein precursor (GPC) gene mediate interaction with host cells, yet the organization and functions of the YEZV glycoproteins remain largely undefined. To characterize YEZV glycoproteins in the context of infection, we developed a reverse genetics system that enables recovery of recombinant YEZV entirely from cloned cDNAs. The recombinant wild-type virus exhibited growth properties comparable to those of the parental isolate *in vitro* and maintained pathogenicity *in vivo*. Proteomic analysis of purified virus particles produced in mammalian cells showed that peptide coverage of GPC-derived products began at residue 69. We therefore used the reverse genetics system to examine the functional importance of the GPC subregion upstream of residue 69. A mutant with a deletion of GPC residues 28 to 68 (rYEZV GPCΔ28–68), which retained the predicted signal peptide, was successfully recovered, suggesting that this subregion is dispensable for producing infectious virus. rYEZV GPCΔ28–68 propagated at levels comparable to those of the wild-type virus in mammalian cells and exhibited similar pathogenicity in a mouse model. Interestingly, the mutant reached lower viral titers in tick-derived ISE6 cells and in ticks *in vivo*, suggesting that this N-terminal subregion may contribute to virus replication in ticks. Together, these findings demonstrate the utility of the newly established reverse genetics system for dissecting the functions of the YEZV glycoproteins in mammalian and tick systems.

**Author Summary:** Viral glycoproteins are important structural components on the surface of enveloped viruses and enable viruses to attach to and enter host cells. In orthonairoviruses, two glycoproteins Gn and Gc are produced from a single glycoprotein precursor (GPC) gene. However, the organization and functions of the glycoproteins of Yezo virus (YEZV), an emerging tick-borne orthonairovirus associated with febrile disease in East Asia, remain poorly understood. To characterize the YEZV glycoproteins, we developed a reverse genetics system for YEZV that allows us to engineer recombinant viruses. Using this platform, we found that a subregion spanning residues 28–68 of GPC was dispensable for recovery of infectious virus. In particular, the mutant grew at levels comparable to those of the wild-type virus in mammalian cells and caused similar disease progression in a mouse model but grew at lower levels in tick cells and in ticks. This contrast suggests that the deleted region may contribute to YEZV growth in tick systems. Together, our findings implicate a previously uncharacterized GPC region in host-dependent differences in YEZV replication and demonstrate the utility of reverse genetics for functional analysis of YEZV proteins.

## Introduction

The genus *Orthonairovirus* within the family *Nairoviridae* comprises a large and diverse group of bunyaviruses, including medically and veterinarily important pathogens such as Crimean-Congo hemorrhagic fever virus (CCHFV) [1], Nairobi sheep disease (NSD) virus [2], Dugbe virus [3], and Tamdy virus [4]. Many orthonairoviruses are tick-associated or tick-borne [5], and well-characterized pathogenic members such as CCHFV are maintained in tick-vertebrate transmission cycles [6,7]. In recent years, several emerging orthonairoviruses, including Yezo virus (YEZV) [8,9], Songling virus [10], Beiji nairovirus [11], and Wetland virus [12], have been associated with human disease. YEZV belongs to the Sulina genogroup and was first identified in human patients with a history of tick bites [8]. Following its initial identification in Hokkaido, Japan, several cases have been identified in Northeastern China [9,13,14] and Russia [15]. Together with other emerging tick-borne orthonairoviruses within the Sulina and Tamdy genogroups, YEZV has become a public health concern in East Asia.

Orthonairoviruses possess tri-segmented RNA genomes, S, M, and L. The M segment encodes a glycoprotein precursor (GPC) that is processed into two structural glycoproteins: Gn and Gc. GPC processing in orthonairoviruses is complex and is best characterized in CCHFV. In CCHFV, GPC is cleaved by host signal peptidase to produce precursors of Gn and Gc. Gn precursor subsequently undergoes further proteolytic processing by furin and subtilisin kexin isozyme-1/site-1 protease (SKI-1/S1P), generating mature Gn and additional glycoprotein products [16–18]. In addition to the structural glycoproteins, Gn and Gc, glycoprotein products derived from the region upstream of mature Gn, including GP38, have been identified in CCHFV-infected cells [18]. Recent studies suggest that this GP38 may contribute to glycoprotein maturation and infectious particle formation [16,19]. However, other than CCHFV, the detailed processing of GPC into mature glycoproteins remains poorly understood.

Many orthonairoviruses, including YEZV, are associated with tick vectors [5–7,20] and thus need to be maintained efficiently in both vertebrate hosts and arthropod vectors. Among bunyaviruses, specific viral proteins have been shown to contribute to replication or dissemination within the vector. For example, a nonstructural protein, NSm, of mosquito-borne bunyaviruses, i.e., Rift Valley fever virus in the genus *Phlebovirus* and Bunyamwera virus in the genus *Orthobunyavirus*, promotes viral dissemination from the mosquito midgut [21,22]. Another nonstructural protein, NSs, expressed by a tick-borne bandavirus, severe fever with thrombocytopenia syndrome virus, has been demonstrated to suppress antiviral RNA silencing in ticks [23]. While virus-host molecular interactions are necessary for efficiently establishing infections, so far, the viral factors that facilitate adaptation of orthonairoviruses to their tick vectors have not been reported.

In this study, we established a cDNA-clone-based reverse genetics system for YEZV for the first time among the emerging tick-borne orthonairoviruses. We applied the established system to characterize YEZV glycoproteins and found that the N-terminal-most portion before residue 69 is not incorporated into virions and not necessary for YEZV rescue in mammalian cells. Interestingly, the mutant lacking the subregion exhibited reduced viral titers in tick cells and ticks, suggesting that this nonstructural region of YEZV contributes to efficient viral replication within the tick host.

## Results

### Recovery of rYEZV from cDNA clones

Based on existing reverse genetics systems for other orthonairoviruses [16,24,25], we prepared T7 promoter-driven plasmids to produce antigenomic RNAs corresponding to the YEZV L, M, and S segments, with an HDV ribozyme sequence on their 3’ termini (namely pMK-YEZV L, pMK-YEZV M, and pMK-YEZV S(C353A)), and helper plasmids expressing YEZV L and N proteins (namely pCAG-YEZV L and pCAG-YEZV N). These plasmids were transfected into Vero E6 cells together with a plasmid expressing T7 RNA polymerase (pCMV-T7 RNA POL). At 21 to 28 days post-transfection, supernatants were blindly passaged first in *Ixodes scapularis* tick embryo-derived ISE6 cells and then in Vero E6 cells to obtain rYEZV (rYEZV WT; Fig 1A). Recovery of rYEZV was confirmed by the expression of YEZV antigens in cells infected with rYEZV (Fig 1B). To exclude contamination by the parental isolate, a silent mutation (C353A) artificially introduced into the S segment as a genetic marker was confirmed in the rescued virus (Fig 1C).

**Fig 1.**
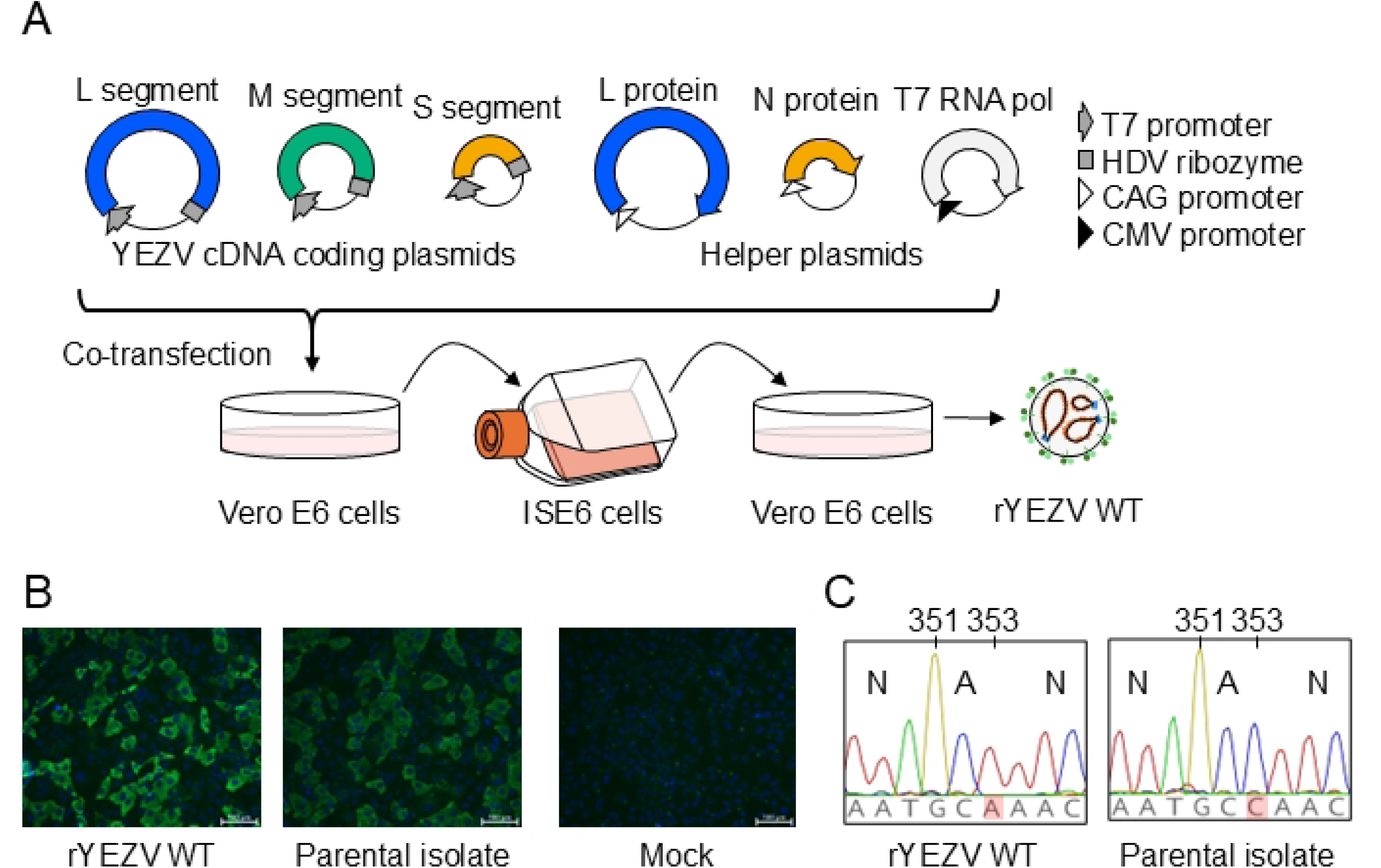
Recovery of a recombinant Yezo virus (rYEZV). (A) Schematic representation of the plasmid-based reverse genetics system workflow used for the recovery of wild-type rYEZV (rYEZV WT). Six plasmids, including three YEZV cDNA clones, were co-transfected into Vero E6 cells. Supernatants were then blindly passaged to ISE6 cells followed by a second blind passage in Vero E6 cells. (B) Immunofluorescence assay of Vero E6 cells inoculated with rYEZV WT or the parental isolate. YEZV antigen is shown in green, and nuclei are counterstained with DAPI (blue). (C) Representative chromatograms showing the sequences of the parental isolate and rYEZV WT at nucleotide position 353 of the S segment to confirm the genetic marker introduced in the corresponding cDNA clone.

### Characterization of rYEZV in vitro and in vivo

To determine whether rescued rYEZV WT retained the growth properties of the parental isolate, we compared the multistep growth kinetics of rYEZV WT and the parental isolate in Vero E6, Hep3B, and ISE6 cells. In all three cell lines tested, rYEZV WT exhibited growth kinetics similar to the parental virus, reaching peak titers at five or seven days post-inoculation (Fig 2A). In Vero E6 cells, the sizes of foci of rYEZV WT were comparable to those of the parental virus (Fig 2B).

**Fig 2.**
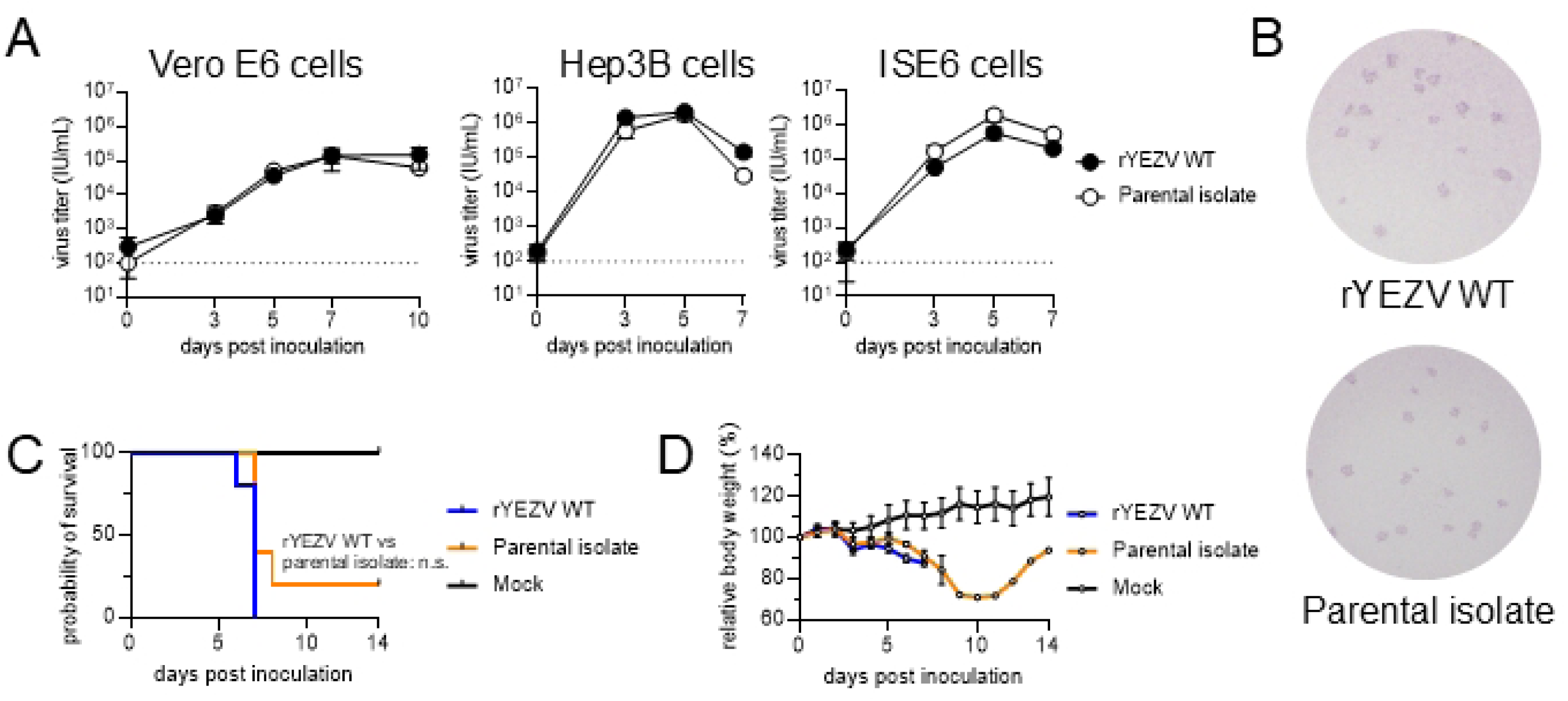
*In vitro* growth properties and *in vivo* pathogenicity of rYEZV. (A) Growth kinetics of rYEZV WT and the parental isolate in Vero E6, Hep3B, and ISE6 cells. Cells were infected at an MOI of 0.001 and supernatants were collected at the indicated time points for virus titration. Broken lines indicate the limit of detection (LOD). Data are shown as the mean ± SD from three replicates. (B) Representative images of focus formation by rYEZV WT and the parental isolate in Vero E6 cells. Infected cells were fixed and stained at seven days post-inoculation with anti-YEZV polyclonal antibodies. (C) Survival curves and (D) relative body weight of mice challenged with rYEZV WT or the parental isolate. AG129 mice (n = 5) were intraperitoneally inoculated with 10^3^ IU/100 µL of rYEZV WT or the parental isolate. The mock group received 100 µL of DMEM. Survival was compared between the rYEZV WT and parental isolate groups using the log-rank test (n.s., not significant). Relative body weight is shown as the mean ± SD.

To assess the pathogenicity of rYEZV WT, we used AG129 mice, which lack type I and type II interferon receptors and were previously established as a lethal model of YEZV infection [26]. The mice were inoculated with either rYEZV WT or parental isolate. Mice infected with rYEZV WT showed survival comparable to that of mice infected with the parental virus (*P* = 0.0926; Fig 2C), and body weight changes were also largely similar between the two virus- infected groups (Fig 2D). Together, these results indicate that rYEZV WT retained the characteristics of the parental virus both *in vitro* and *in vivo*.

### Proteomic and glycoproteomic analysis of YEZV particles

To obtain insights into the organization of glycoproteins on YEZV particles, we performed mass spectrometry-based glycoproteomic and proteomic analyses on purified viral particles. YEZV particles were purified through iodixanol gradient density centrifugation, and the sample quality was evaluated by silver staining following SDS-PAGE. Three major bands with apparent molecular masses of approximately 70, 55, and 40 kDa were observed by silver staining. Immunoblotting with antibodies specific for YEZV proteins showed corresponding bands at similar positions, supporting their assignment as Gc, N, and Gn (Fig 3A). After confirming the successful enrichment of YEZV particles, the purified sample was then subjected to trypsin digestion and subsequent liquid chromatography-tandem mass spectrometry (LC-MS/MS) analysis. Glycoproteomic analysis identified an *O*-glycosylated peptide spanning residues 69–81, which contained residues predicted to undergo *O*- glycosylation by NetOGlyc 4.0 (Fig 3B, S1, and S2 Tables). *N*-linked glycosylation sites at Asn112 (NLS), Asn401 (NGT), Asn463 (NDT), Asn731 (NHS), and Asn1250 (NST) were also identified (Fig 3B and S3–S5 Tables). Proteomic analysis further identified peptides distributed across the region upstream of the putative SKI-1/S1P cleavage motif (RKLL), but the N-terminal-most identified peptide began at residue 69 (Fig 3B and S6 Table). No peptides derived from residues 1–68 were identified in the sample analyzed, suggesting that the N-terminal residues following the putative signal peptide are cleaved at an undefined position upstream of residue 69 and not incorporated into virions.

**Fig 3.**
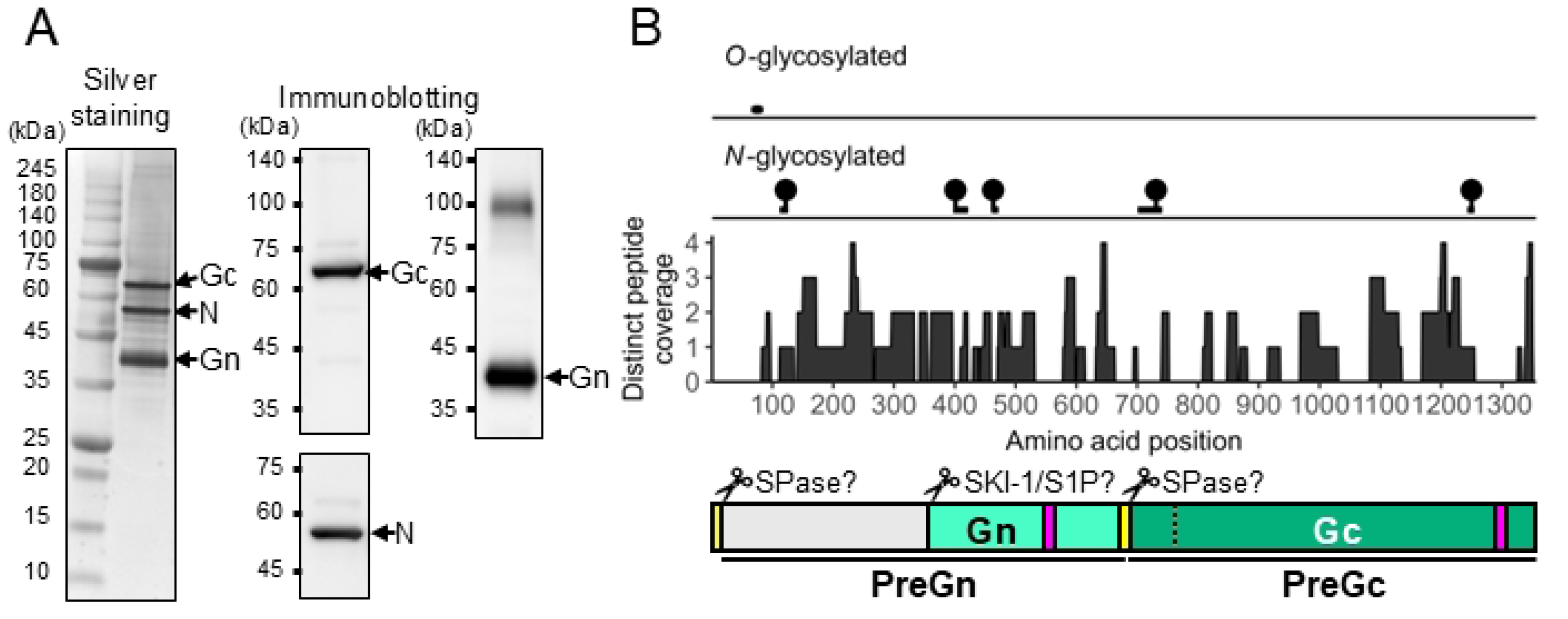
Proteomic and glycoproteomic analysis of purified YEZV particles. (A) Silver staining and immunoblotting of purified YEZV particles under reducing conditions. Immunoblotting was performed using specific antibodies targeting Gn, N, and Gc, respectively. Major protein bands in the silver-stained gel corresponding to those detected by immunoblotting were then identified as YEZV proteins based on their molecular weights. (B) Mapping of glycosylated peptides and proteomic peptide coverage onto the YEZV glycoprotein precursor (GPC). The upper tracks show the identified *O*-glycosylated peptide and *N*- glycosylation sites, followed by distinct peptide coverage across the GPC sequence. The lower schematic shows the predicted signal peptide (yellow), predicted transmembrane regions (magenta), putative mature Gn and Gc regions, and putative cleavage sites indicated by scissors. Predicted signal peptide and transmembrane regions are shown based on analyses using SignalP 6.0, SOSUI, and TMHMM 2.0 (S1–S3 Figs). Regions of precursors of Gn and Gc (PreGn and PreGc, respectively) are indicated below the schematic. The unidentified cleavage site upstream of mature Gn is indicated by a broken line.

### Processing of the YEZV GPC region upstream of mature Gn

We next examined the molecular fate of the N-terminal region upstream of residue 69, from which no peptides were identified by proteomic analysis. To track this region, we constructed GPC expression plasmids containing a peptide tag immediately after residue 41, 345, or 367, hereafter referred to as GPC[FLAG41], GPC[FLAG345], or GPC[FLAG367], respectively.

GPC[FLAG345] and GPC[FLAG367] were designed to flank the putative SKI-1/S1P cleavage motif (RKLL; residues 354–357). Regardless of the position where the peptide tag was inserted, immature proteins such as full-length GPC and PreGn (defined as the uncleaved Gn precursor) were detected by using anti-FLAG antibodies. Mature Gn and Gc were also expressed in all cells (Fig 4B–D). However, the FLAG-tagged mature Gn was identified only in GPC[FLAG367]-expressing cells, supporting cleavage at or near the putative SKI-1/S1P cleavage motif (Fig 4B). No discrete fragment of approximately 5 kDa, which would be expected if cleavage occurred near residue 68, was detected in either cell lysates or culture supernatants of cells expressing GPC[FLAG41] (data not shown). Together, these findings suggest that the N-terminal-most region of YEZV GPC may undergo distinct processing, although its precise molecular status remains unresolved.

**Fig 4.**
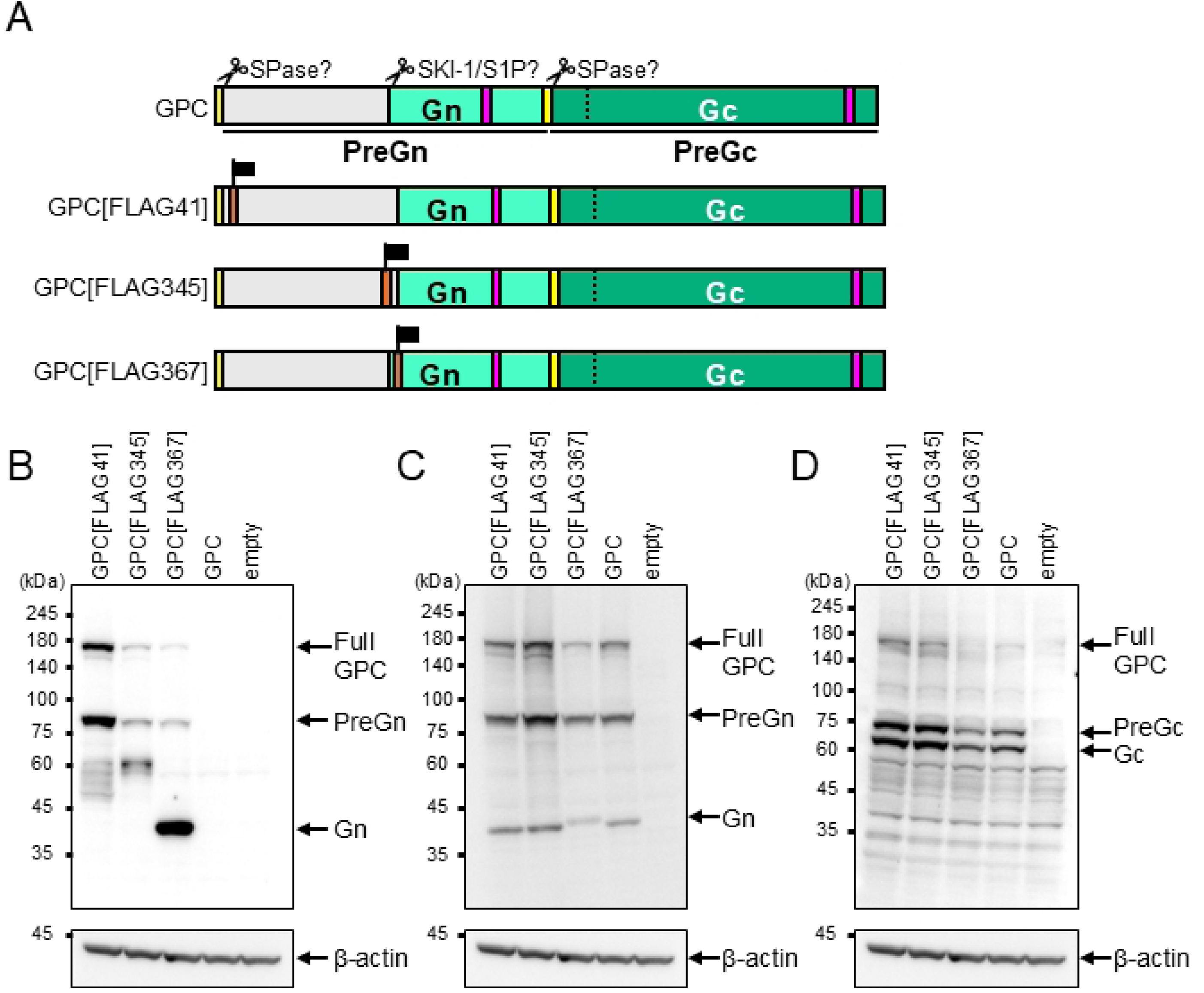
Characterization of the N-terminal region upstream of mature Gn. (A) Schematic representation of YEZV GPC expression constructs containing a FLAG tag (DYKDDDDK) at residue 41, 345, or 367. FLAG insertion sites are indicated by orange boxes with flags. The predicted signal peptide and transmembrane region are shown in yellow and magenta, respectively. (B–D) Immunoblotting of lysates from HEK293T cells transfected with the YEZV GPC expression constructs. The top panels show membranes probed with anti-FLAG (B), anti-Gn (C), and anti-Gc (D) antibodies, and the bottom panels show membranes probed with anti-β-actin antibodies.

### Rescue of a mutant YEZV lacking GPC residues 28–68

Because the N-terminal 68 residues of GPC were absent from the virions and their fate remained unclear, we next used the reverse genetics system to directly test whether this region is required for recovery of infectious virus. We therefore generated an M-segment clone lacking amino acids 28–68 within the GPC open reading frame, designated pMK-YEZV MΔ28–68. In this construct, the predicted signal peptide was intentionally retained based on the proposed GPC organization of orthonairovirus glycoprotein precursors (Fig 5A). Using the same reverse genetics procedure as for rYEZV WT, pMK-YEZV M was replaced with pMK-YEZV MΔ28– 68 and transfected into Vero E6 cells together with the other plasmids. After blind passage of culture supernatants, viral antigen was detected by immunofluorescence assay in Vero E6 cells inoculated with rYEZV GPCΔ28–68 (Fig 5B), and the mutant and wild-type viruses had grossly similar particle morphologies (Fig 5C). We next examined whether the deletion affected viral protein expression, especially GPC processing. N, mature Gn, and mature Gc were detected in cells infected with both viruses, and PreGn of rYEZV GPCΔ28–68 showed the expected mobility shift to a lower molecular mass resulting from the deletion (Fig 5D). In the purified viral particles, N, Gn, and Gc were detected in both viruses, with no obvious differences in the apparent molecular masses of the major bands (Fig 5E), except for the incorporation of a PreGn- like protein in rYEZV GPCΔ28–68 particles. Together, these results indicate that deletion of GPC residues 28–68 did not impair the formation of particles containing the major structural proteins, although the maturation of Gn may have been affected. Since the N-terminal amino acid sequence was highly conserved among YEZV strains, but not among viruses in the Sulina genogroup (Fig 5F), the removal of this subregion from mature Gn is expected to be YEZV- specific.

**Fig 5.**
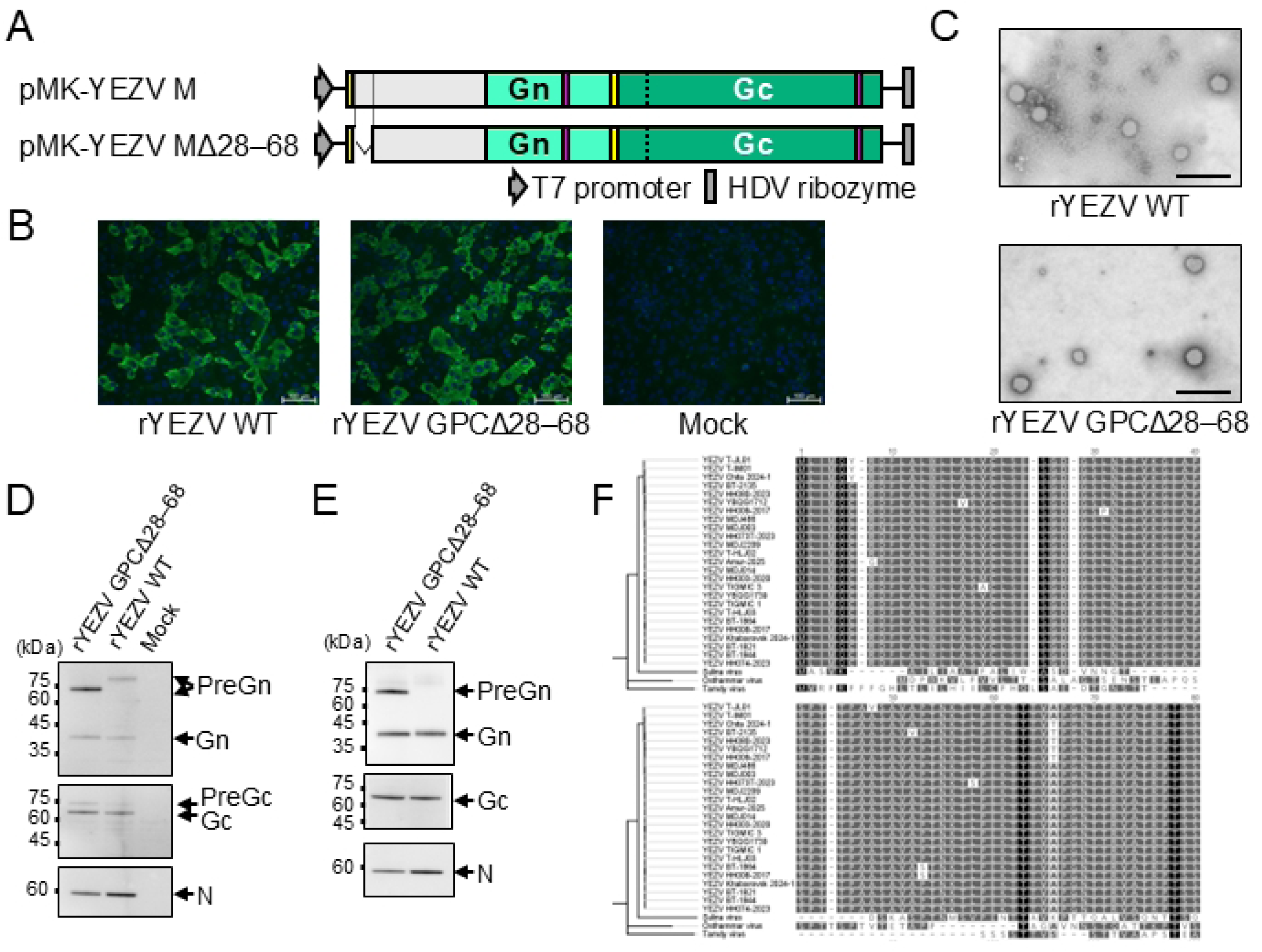
Rescue of recombinant YEZV mutant lacking GPC residues 28–68. (A) Schematic representation of the wild-type (pMK-YEZV M) and mutant (pMK-YEZV MΔ28–68) M-segment plasmids used for recovery of recombinant YEZV. (B) Immunofluorescence assay of Vero E6 cells inoculated with rYEZV GPCΔ28–68, rYEZV WT, or DMEM (mock). YEZV antigen is shown in green, and nuclei are counterstained in blue with DAPI. (C) Transmission electron microscopy of purified virions of rYEZV WT and rYEZV GPCΔ28–68. Scale bars, 500 nm. (D and E) Immunoblotting of lysates from Vero E6 cells infected with rYEZV GPCΔ28–68 or rYEZV WT, or mock-infected cells (D), and purified virions from culture supernatants (E), using anti-Gn (top), anti-Gc (middle), and anti-N (bottom) antibodies. (F) Amino acid sequence comparison of the N-terminal region of GPC from 24 strains of YEZV, Östhammar virus, Sulina virus, and Tamdy virus. A multiple sequence alignment corresponding to GPC residues 1–80 is shown. A neighbor-joining tree based on GPC residues 1–100 is shown to the left of the alignment.

### Growth and pathogenicity of rYEZV GPCΔ28–68 in mammalian systems

First, we investigated rYEZV GPCΔ28–68 in mammalian cells and the mouse model. We compared multistep growth kinetics of the recombinant viruses in Vero E6 and Hep3B cells. Cells were infected with rYEZV WT or rYEZV GPCΔ28–68 at an MOI of 0.001. In Vero E6 and Hep3B cells, rYEZV GPCΔ28–68 reached peak titers earlier than rYEZV WT (Fig 6A). In AG129 mice, the two viruses exhibited no significant difference in survival curves (*P* = 0.0926; Fig 6B) and caused broadly similar body weight changes (Fig 6C). These findings indicate that deletion of GPC residues 28–68 altered viral growth kinetics in mammalian cells but did not markedly reduce peak titers in these cells or attenuate pathogenicity in the mouse model.

**Fig 6.**
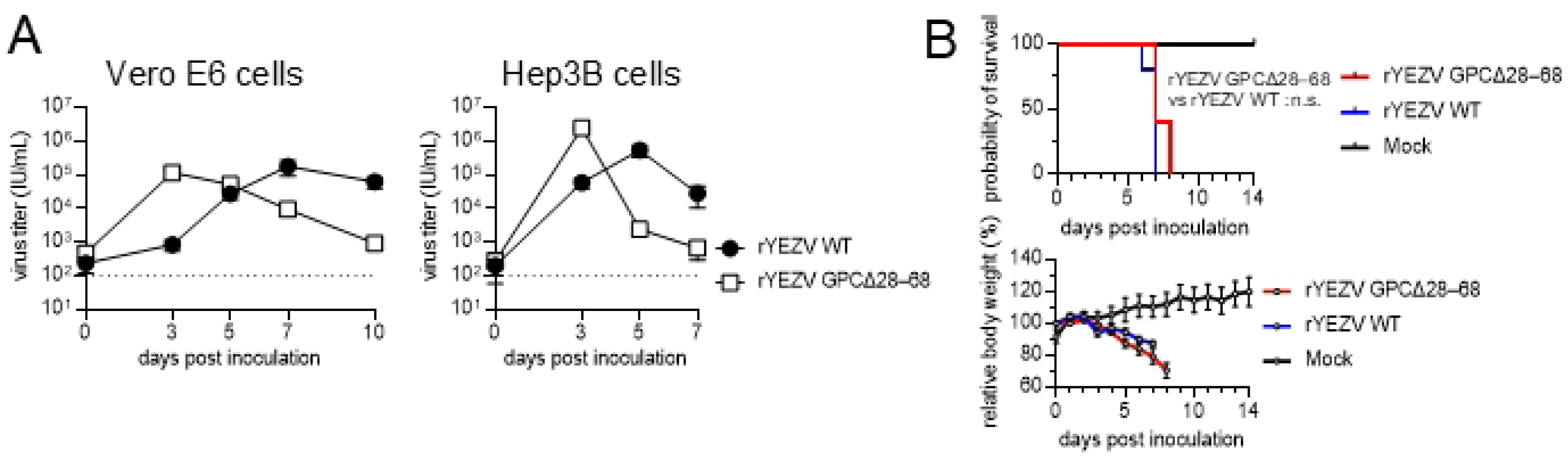
Characterization of rYEZV GPCΔ28–68 *in vitro* and *in vivo* in mammalian systems. (A) Growth kinetics of rYEZV GPCΔ28–68 and rYEZV WT in Vero E6 and Hep3B cells. Cells were infected at an MOI of 0.001, and culture supernatants were collected at the indicated time points for virus titration. Broken lines indicate the limit of detection (LOD). Data are shown as the mean ± SD from three replicates. (B) Survival curves and (C) relative body weight of AG129 mice challenged with rYEZVs. AG129 mice (n = 5) were intraperitoneally inoculated with 10^3^ IU/100 µL of rYEZV GPCΔ28–68 or rYEZV WT. The negative control group received 100 µL of DMEM. Survival was compared between the rYEZV GPCΔ28–68 and rYEZV WT groups using the log-rank test (n.s., not significant). Relative body weight is shown as the mean ± SD.

### Replication of rYEZV GPCΔ28–68 in the tick host

Since deletion of GPC residues 28–68 did not substantially impair viral growth in mammalian cells or pathogenicity in the mouse model, we next evaluated viral replication in tick systems. *Ixodes scapularis* embryo-derived ISE6 and *Ixodes ricinus* embryo-derived IRE/CTVM20 cells were inoculated with either rYEZV WT or rYEZV GPCΔ28–68 at an MOI of 0.001. In ISE6 cells, rYEZV GPCΔ28–68 reached a lower peak titer than rYEZV WT (Fig 7A), while in IRE/CTVM20 cells, rYEZV GPCΔ28–68 reached its peak earlier but attained a peak titer comparable to that of rYEZV WT, similar to the growth observed in Vero E6 and Hep3B cells. Given these cell-type-dependent phenotypes in tick-derived cells, we next examined viral replication in *Ixodes persulcatus* and *Ixodes ovatus*, two tick species associated with YEZV in nature [8,9,20,27]. Adult females of both species were microinjected with either rYEZV WT or rYEZV GPCΔ28–68, and virus titers were measured at seven days post-inoculation. In both tick species, titers of rYEZV GPCΔ28–68 were significantly lower than those of rYEZV WT (Fig 7B). These findings indicate that the deletion of GPC residues 28–68 compromises viral fitness in the tick host.

**Fig 7.**
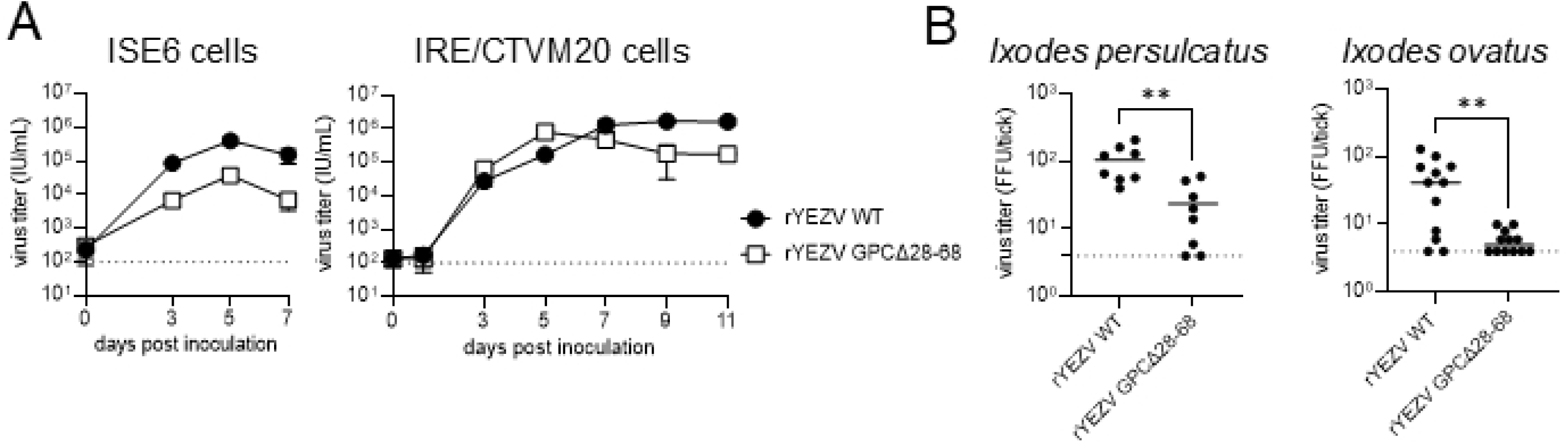
Characterization of rYEZV GPCΔ28–68 *in vitro* and *in vivo* in tick systems. (A) Growth kinetics of rYEZV GPCΔ28–68 and rYEZV WT in ISE6 and IRE/CTVM20 cells. Cells were infected at an MOI of 0.001, and culture supernatants were collected at the indicated time points for virus titration. Data are shown as the mean ± SD from three replicates. (B) Virus titers of *Ixodes persulcatus* and *Ixodes ovatus* ticks inoculated with rYEZV GPCΔ28–68 or rYEZV WT by microinjection. Virus titers of the two viruses were compared using the Mann– Whitney U test (\*\**P* < 0.01). Broken lines indicate the limit of detection (LOD).

## Discussion

YEZV is one of the emerging tick-borne orthonairoviruses associated with febrile illness in East Asia, yet its molecular characteristics remain poorly understood. In this study, we newly established a reverse genetics system to investigate an N-terminal subregion of the YEZV GPC whose absence was suggested by proteomic analysis of virus particles. Deletion of this subregion had no substantial effects in mammalian systems but reduced viral replication in tick-derived cells and in ticks. Thus, our findings identify a previously uncharacterized region of the YEZV GPC that may contribute to efficient viral replication in tick environments.

Reverse genetics systems have been reported for only a limited number of orthonairoviruses, including CCHFV, Hazara virus, and Tofla virus [16,24,25]. YEZV and emerging viruses belonging to the Sulina and Tamdy genogroups are phylogenetically distant from these well-studied orthonairoviruses [8,9]. Thus, to our knowledge, this study provides the first reverse genetics platform for these Sulina and Tamdy orthonairoviruses recently recognized as human pathogens. Given the substantial diversity within the genus *Orthonairovirus*, this system expands the landscape of molecular studies of emerging members of the genus. Nevertheless, the relatively long time required for virus recovery leaves room for further optimization of the system.

Among orthonairoviruses, the glycoprotein undergoes complex proteolytic processing, and thus, clarifying the actual forms of mature glycoproteins on viral particles is crucial for a molecular-level understanding of their functions. Our findings suggest that the overall organization and processing of YEZV GPC are broadly consistent with those described for other orthonairoviruses. Our data support cleavage at or near the putative SKI-1/S1P cleavage motif, and the GP38-like protein upstream of mature Gn is also found on the virus particle, as reported for CCHFV GP38 [19,28,29]. Interestingly, our study demonstrated that the region further upstream of the GP38-like protein (residues 1–68) was nonstructural and was not incorporated into YEZV particles. Due to the possibility of multiple cleavages within this region, including one by a signal peptide peptidase, we could not determine the fate of the nonstructural product(s) in the present study. Thus, while YEZV appears to share major features of GPC processing with other orthonairoviruses, the precise processing and molecular status of its N-terminal region remain to be clarified.

The successful recovery of rYEZV GPCΔ28–68 indicates that this region is not essential for the production of infectious virus, and the deletion did not alter the overall profile of the major structural proteins, except for the incorporation of a PreGn-like protein, suggesting that the deletion may affect GPC processing. Despite these changes, deletion of residues 28–68 did not substantially impair viral growth in mammalian cells or attenuate pathogenicity in the mouse model, indicating that this region is largely dispensable for viral replication in mammalian systems. In contrast, rYEZV GPCΔ28–68 exhibited reduced viral titers in ISE6 cells and in experimentally infected ticks but not in IRE/CTVM20 cells. This host- or cell-type-dependent phenotype raises the possibility that this region is involved in adaptation to particular tick hosts, potentially reflecting interactions with species-specific host factors. However, it remains unclear whether this phenotype reflects a specific function of residues 28–68 as a nonstructural protein or indirect effects of the deletion on GPC processing, particle composition, or genome organization.

In conclusion, this study describes a reverse genetics system for YEZV and provides a platform for molecular analysis of this emerging orthonairovirus. Using this technique, we obtained initial evidence that GPC residues 28–68, corresponding to the N-terminal-most subregion examined in this study, are dispensable for recovery of infectious virus. The precise molecular state of this subregion remains to be defined, including its role in tick-derived cells and in the tick host. Thus, the present work not only expands the range of orthonairoviruses for which genetic tools are available but also provides a practical framework for future studies of YEZV glycoprotein biology and host adaptation.

## Materials and Methods

### Cells and Viruses

Vero E6 (African green monkey kidney; JCRB9007, JCRB), Hep3B (human liver carcinoma; HB-8064, ATCC), and HEK293T (human embryonic kidney) cells were cultured in Dulbecco’s Modified Eagle’s Medium (DMEM, Nacalai Tesque) supplemented with 10% FBS (ICN Biomedicals), 1% penicillin-streptomycin (Thermo Fisher Scientific), and 1% L-glutamine (FUJIFILM Wako Chemicals) at 37°C with 5% CO_2_. ISE6 (*Ixodes scapularis* embryo) cells were kindly provided by the CEH Institute of Virology and Environmental Microbiology (Oxford, UK). ISE6 cells were cultured in Leibovitz L-15B medium supplemented with 10% FBS and 5% tryptose phosphate broth (Sigma-Aldrich) at 32°C. IRE/CTVM20 (*Ixodes ricinus* embryo) cells were cultured in Leibovitz L-15 mixed with L- 15B medium, supplemented with 10% tryptose phosphate broth, 10% FBS, 1% penicillin-streptomycin, 1% L-glutamine, and 0.1% bovine lipoprotein concentrate (MP Biomedicals) at 28°C.

YEZV strain HH003-2020 was isolated from patients as previously reported [8]. Briefly, the virus was isolated by inoculating Vero E6 cells with serum collected from AG129 mice that had been injected with plasma from an infected patient. All experiments with infectious materials were conducted in a biosafety level 2 (BSL-2) facility at the International Institute for Zoonosis Control, Hokkaido University.

### Plasmids

YEZV RNA was extracted using NucleoSpin RNA Virus (TaKaRa) according to the manufacturer’s instructions. YEZV cDNA was synthesized from viral RNA using PrimeScript II (TaKaRa) with either random primers or a gene-specific primer complementary to the 3′ end of the viral genomic RNAs. The open reading frame (ORF) of each segment was amplified using KOD One (TOYOBO) and cloned into the pCAGGS vector to generate the expression plasmids: pCAG-YEZV L, pCAG-YEZV GPC, and pCAG- YEZV N. pCMV-T7 RNA POL was a gift from Baisong Lu (Addgene plasmid #138352; http://n2t.net/addgene:138352; RRID:Addgene_138352).

For construction of full-length clones, a pMK backbone containing the T7 promoter, untranslated regions (UTRs) of each YEZV segment RNA, and a hepatitis delta virus (HDV) ribozyme sequence was first assembled by PCR amplification using KOD One (TOYOBO), followed by In-Fusion cloning (TaKaRa) according to the manufacturer’s instructions. ORF fragments derived from the expression plasmids were then amplified using KOD One (TOYOBO) or PrimeSTAR Max (TaKaRa) and inserted into this vector by In-Fusion cloning to generate the full-length YEZV plasmids pMK-YEZV L, pMK-YEZV M, and pMK-YEZV S of antigenomic sense. Then, a genetic marker was introduced into the pMK-YEZV S by site-directed mutagenesis to produce pMK-YEZV S(C353A). A plasmid encoding GPC lacking amino acid residues 28–68 (pMK-YEZV MΔ28–68) was generated from pMK-YEZV M by site-directed mutagenesis. Briefly, PCR amplification was performed using PrimeSTAR Max (TaKaRa) and primers to introduce the desired mutation or deletion, followed by phosphorylation with T4 polynucleotide kinase (New England Biolabs) and ligation with T4 DNA ligase (New England Biolabs).

Expression plasmids encoding GPC with a FLAG tag (DYKDDDDK) inserted at various positions, i.e., pCAG-YEZV GPC[FLAG41], pCAG-YEZV GPC[FLAG345], and pCAG-YEZV GPC[FLAG367], were generated from pCAG-YEZV GPC by site-directed mutagenesis using primers containing the FLAG-tag insertion sequence. PCR amplification was performed using KOD Fx Neo (TOYOBO), followed by phosphorylation with T4 polynucleotide kinase (New England Biolabs) and ligation with T4 DNA ligase (New England Biolabs). Primers used for this study are listed in S7 Table.

### Rescue of recombinant YEZVs (rYEZVs) from plasmid DNAs

Vero E6 cells were seeded in six-well plates at 5.0 × 10^5^ cells/well in 2 mL complete medium, 16–18 h prior to transfection. Cells were then transfected with 1 μg each of pMK-YEZV L, pMK-YEZV M (or pMK-YEZV MΔ28–68), and pMK-YEZV S(C353A), together with 0.67 μg of pCAG-YEZV N, 0.33 μg of pCAG-YEZV L, and 1 μg of pCMV-T7 RNA POL, using Lipofectamine 3000 and P3000 reagent (Invitrogen) at 3 μL and 2 μL per 1 μg of DNA, respectively, in a total volume of 250 μL Opti-MEM (Gibco). Three days after the transfection, the culture medium was replaced with DMEM containing 2% FBS. The cells were then cultured from 21 to 28 days post-transfection, with the medium replaced once a week with fresh 2% FBS DMEM. Since direct blind passage only in Vero E6 cells did not yield rescue stocks with reproducible growth properties in preliminary experiments, culture supernatants from the transfected cells were first subjected to blind passage in ISE6 cells, followed by a second blind passage in Vero E6 cells. The supernatant of Vero E6 cells was used as the recombinant virus stock and was stored at –80°C until use.

### Virus titration

For immunofluorescence-based infectivity assay, Vero E6 cells were seeded in 96-well plates at 2.5 × 10^4^ cells/well and cultured for 24 h. The cells were then inoculated with 10-fold serial dilutions of the virus prepared in DMEM containing 2% FBS. At 24 h post-infection, the cells were fixed with 4% paraformaldehyde (PFA; FUJIFILM Wako Chemicals) in phosphate-buffered saline (PBS). The cells were then stained with heat-inactivated sera from YEZV- infected mice diluted in dilution buffer (0.3% Triton X-100 and 3% bovine serum albumin in PBS), followed by goat anti-mouse IgG conjugated to Alexa Fluor Plus 594 (Invitrogen) diluted in PBS. After staining, images were acquired using an IN Cell Analyzer (Cytiva), and the number of YEZV antigen-positive cells was automatically quantified using IN Cell Developer Toolbox software 1.9.2 (Cytiva). Virus titers were calculated from the number of antigen-positive cells and the corresponding dilution factor and expressed as infectious units per milliliter (IU/mL).

For focus-forming assay, Vero E6 cells were seeded in 24-well plates at 1.3 × 10^5^ cells/well and cultured for 24 h. Samples diluted in serum-free DMEM were inoculated into Vero E6 cells and incubated at 37°C for 1 h. The inoculum was aspirated, and the cells were washed once with PBS. Treated cells were cultured in Eagle’s minimum essential medium (Nissui Pharmaceutical) containing 1% methylcellulose, 1% penicillin-streptomycin, 1% L-glutamine, and 2% FBS for six days. The cells were then fixed with 4% PFA and stained with heat-inactivated sera of YEZV-infected mice diluted in dilution buffer (as described above), followed by incubation with goat anti-mouse IgG H&L-HRP (Abcam). YEZV-infected cell foci were visualized by ImmPACT VIP HRP substrate (Vector Laboratories). Images were obtained by an ImmunoSpot analyzer (Cellular Technology Limited). In the tick infection experiment described below, focus-forming units (FFU), determined from the number of foci in each well, were used to titrate YEZV grown in infected ticks.

### Immunofluorescence assay and sequencing analysis

To confirm the expression of virus antigen in the cells infected with YEZV, Vero E6 cells were inoculated with recombinant viruses or the parental virus. After 24 h of incubation, the cells were fixed with 4% PFA in PBS and stained with heat-inactivated sera of YEZV- infected mice diluted in the dilution buffer (as described above), followed by visualization with Alexa Fluor 488-conjugated goat anti-mouse IgG (Invitrogen). Nuclei were counterstained with DAPI. Fluorescence images were acquired using a fluorescence microscope (Zeiss). Images were merged in ImageJ [30].

To determine the sequences of the recombinant viruses, viral RNAs were extracted by NucleoSpin RNA Virus (TaKaRa), then reverse-transcribed using PrimeScript II (TaKaRa), according to the manufacturer’s protocol. Each segment was then amplified with KOD One (TOYOBO). Nucleotide sequences of the PCR products were determined using BigDye Terminator v3.1 Cycle Sequencing Kit (Thermo Fisher Scientific) and a 3500xL Genetic Analyzer (Applied Biosystems). Sequences were analyzed and visualized by Geneious Prime 2026.1.2 (Dotmatics).

### Growth kinetics

Vero E6 (1.3 × 10^5^ cells/well), Hep3B (1.5 × 10^5^ cells/well), and ISE6 cells (2.5 × 10^5^ cells/well) were seeded in 24-well plates, and IRE/CTVM20 cells (1.0 × 10^6^ cells/well) were seeded in a 6-well plate. All cells were cultured for 24 h prior to infection. For ISE6 and IRE/CTVM20 cells, plates coated with ε-poly-L-lysine (Cosmo Bio) were used. The cells were infected at a multiplicity of infection (MOI) of 0.001. After adsorption for 1 h (37°C for Vero E6 and Hep3B cells, 32°C for ISE6 cells, 28°C for IRE/CTVM20 cells), the inoculum was removed and replaced with fresh medium after washing once with PBS. Vero E6 and Hep3B cells were maintained in 2% FBS DMEM, while ISE6 and IRE/CTVM20 cells were maintained in their corresponding maintenance media. Culture supernatants were collected at the indicated time points, and virus titers were determined as described above.

### Mouse infection experiments

AG129 mice, double-knockout immunocompromised mice lacking both type I and type II interferon receptors, were obtained from Marshall BioResources and maintained by in-house breeding. Five four-week-old AG129 mice of mixed sex were intraperitoneally inoculated with 100 μL of DMEM containing 10^3^ IU of YEZV isolate or an rYEZV, or with DMEM alone. All mice were monitored for body weight changes and survival for 14 days.

All animal experiments were conducted in a BSL-2 laboratory at the International Institute for Zoonosis Control, Hokkaido University under ethical approval (Approval number: 23-0063) and were performed according to the committee’s guidelines.

### Immunoblotting and sample preparation

For preparation of samples for immunoblotting, HEK293T cells were seeded in six-well plates at 9.0 × 10^5^ cells/well one day before transfection and transfected with FLAG-tagged GPC plasmids using Lipofectamine 3000 and P3000 reagent (Invitrogen) at 3.75 μL and 3 μL per 1.5 μg of DNA, respectively, in a total volume of 250 μL Opti-MEM (Gibco). Transfected cells were lysed using the Ez RIPA Lysis Kit (ATTO) according to the manufacturer’s protocol one day after transfection. For preparation of lysates from virus-infected cells, Vero E6 cells were seeded in six-well plates at 5.0 × 10^5^ cells/well and cultured for 24 h. Cells were then infected with rYEZV WT or rYEZV GPCΔ28–68 at an MOI of 0.001 and lysed at three to seven days post-infection using the Ez RIPA Lysis Kit (ATTO) according to the manufacturer’s protocol. For preparation of virus samples, Vero E6 cells were infected with each virus at an MOI of 0.001, and culture supernatants were collected at the time of peak virus production. The supernatants were clarified by centrifugation at 3,500 rpm for 10 min at 4°C, layered onto a 16% iodixanol cushion prepared by diluting OptiPrep (Serumwerk) in PBS, and ultracentrifuged in a Beckman SW41 rotor at 28k rpm for 2 h at 4°C. The resulting pellets were resuspended in PBS.

Samples were mixed with an equal volume of EzApply sample buffer with or without DTT (ATTO) and boiled at 95°C for 5 min. SDS-PAGE was performed using a 10–20% e-PAGEL gradient gel (ATTO) at a constant current of 21 mA for 70 min. For immunoblotting, the separated proteins were transferred to a PVDF membrane using a Qblot Kit M (ATTO). The membranes were blocked with 3% skim milk (FUJIFILM Wako Chemicals) in Tris-buffered saline containing Tween 20 (TBS-T; Takara Bio). The membranes were incubated overnight at 4°C with anti-FLAG M2 antibody (Sigma-Aldrich), or rabbit antisera raised against YEZV- N peptide, YEZV-Gn peptide (amino acids 349–369), or YEZV-Gc peptide (amino acids 803–823) (1:1,000; Cosmo Bio) diluted in TBS-T containing 0.5% skim milk. The membranes were then incubated for 1 h at 25°C with goat anti-mouse IgG H&L-HRP or goat anti-rabbit IgG H&L-HRP (1:10,000; Abcam) in TBS-T with 0.5% skim milk. Proteins were detected using Clarity Western ECL Substrate (Bio-Rad) and visualized with a LuminoGraph III (ATTO).

### Proteomic analysis via LC-MS/MS

Vero E6 cells were seeded in T75 flasks at 4.5 × 10^5^ cells/flask and infected with YEZV isolate at an MOI of 0.01. After 1 h of adsorption, 15 mL of 2% FBS DMEM was added. Five to six days post-inoculation, supernatants were harvested and clarified by centrifugation at 3,500 rpm for 10 min at 4°C. Viruses were concentrated by ultrafiltration with Vivaspin Turbo 15 devices (MWCO 100 kDa; Sartorius) by centrifugation at 2000 rcf for 30 min at 20°C. The viruses were further purified by iodixanol density gradient centrifugation using OptiPrep (Serumwerk) at 8% to 40% iodixanol in PBS in a Beckman SW41 rotor at 28k rpm for 2 h at 4°C. The visible band was collected, diluted in PBS, and centrifuged in a Beckman SW31 rotor at 28k rpm for 2 h at 4°C. The virus pellet was resuspended in PBS and incubated at 4°C until complete resuspension. For sample quality confirmation, resuspended viruses were mixed with an equal volume of EzApply sample buffer containing DTT (ATTO) and boiled at 95°C for 5 min. SDS-PAGE was performed using a 10–20% e-PAGEL gradient gel (ATTO) at a constant current of 21 mA for 70 min. Gels were stained with an EzStain Silver kit (ATTO) according to the manufacturer’s instructions after electrophoresis. Images were acquired using a LuminoGraph III (ATTO).

The resuspended virus pellet was then precipitated with acetone (final concentration of 80%) at −20°C for 1 h. After centrifugation at 14000 rcf for 15 min, the precipitates were recovered and dissolved with 100 mM Tris-HCl (pH 9.0) containing 12 mM sodium deoxycholate and 12 mM sodium lauroylsarcosinate. The proteins were reduced with dithiothreitol, alkylated with iodoacetamide, and quenched with dithiothreitol. The samples were then diluted fourfold with 50 mM ammonium bicarbonate and digested with rLys-C endopeptidase (Promega), followed by Trypsin Gold (Promega). An aliquot of each digest was subjected to LC-MS/MS analysis for proteomic profiling, and the acquired MS/MS spectra were searched against a protein sequence database (SwissProt 40,003 sequences) containing YEZV protein sequences using Mascot (Matrix Science). For glycopeptide analysis, another aliquot of each digest was subjected to enrichment using an Amide-80 column (TOSOH), and the obtained glycopeptides were identified by LC-MS/MS analysis, followed by database searching using Byonic (Protein Metrics). In addition, an aliquot of each digest was treated with PNGase F (Takara Bio) in H_2_ ^18^O according to the isotope-coded glycosylation site-specific tagging (IGOT) method for site-specific *N*-glycosylation analysis [31,32]. The resulting peptides were lyophilized, redissolved in nonlabelled water, treated with trypsin at 4°C overnight, and purified using GL-Tip SDB (GL Sciences, Tokyo, Japan). LC-MS/MS analysis was performed using an UltiMate 3000 RSLCnano system coupled to an Orbitrap Eclipse Tribrid mass spectrometer (Thermo Fisher Scientific). Peptides were separated on a C18 tip column (inner diameter: 0.075 mm, length: 25 cm, 1.9 μm particle, Nikkyo Technos, Tokyo, Japan) at a flow rate of 300 nL/min using a linear gradient of 3%–35% acetonitrile in 0.1% formic acid over 45 min. Separated peptides were ionized by electrospray ionization and analyzed in positive-ion mode using data-dependent acquisition. For proteome and IGOT analyses, MS and MS/MS spectra were acquired using the Orbitrap analyzer at resolutions of 60,000 and 30,000, respectively. Selected precursor ions were fragmented by higher-energy collision- induced dissociation (HCD) with a normalized collision energy of 30%. For glycoproteome analysis, MS and MS/MS spectra were acquired using the Orbitrap analyzer at resolutions of 120,000 and 60,000, respectively, and selected precursor ions were fragmented by HCD using stepped normalized collision energies of 20%, 30%, and 40%. Obtained MS/MS spectra from the proteome and IGOT analyses were searched using Mascot (ver. 2.8.0.1), whereas those from the glycoproteome analysis were searched using Byonic (ver. 4.3.4). A Swiss-Prot human FASTA database downloaded on May 19, 2020, containing YEZV protein sequences, was used for database searching. Enzyme cleavage setting was used Trypsin (C-terminus KR) and maximum missed cleavages were allowed 2. Full tryptic specificity was used for the proteome and glycoproteome analyses, whereas both full and semi-tryptic searches were performed for the IGOT analysis. The precursor and fragment mass tolerances were set to 2 and 5 ppm, respectively. Peptide N-terminal pyroglutamine formation (Q), peptide N-terminal ammonia loss (C), oxidation (M), and Delta:H(1) N(-1) 18O(1) (N), the latter only for IGOT analysis, were considered variable modifications. In the Mascot search, peptides with pep_rank = 1 and pep_expect values below the significance threshold were considered identified. For the IGOT analysis, only peptides containing the Delta:H(1) N(-1) 18O(1) modification on an Asn residue within the *N*-glycosylation consensus sequence (sequon; Asn- Xaa-[Ser/Thr/Cys], Xaa ≠ Pro) were selected as IGOT peptides. In the Byonic search, peptides with Confidence = High were considered as “identified”.

#### Sequence comparison and phylogenetic analysis of the N-terminal region of GPC

Amino acid sequences of the GPC from 24 strains of YEZV, Östhammar virus [33], Sulina virus, and Tamdy virus were aligned using MUSCLE in Geneious Prime 2026.1.2 (Dotmatics). A neighbor-joining tree was constructed from the alignment. Amino acid residues 1–80 were used for comparative sequence alignment of the N-terminal region. Accession numbers of sequences used are listed in S8 Table.

### Tick infection by microinjection

Adult female *Ixodes persulcatus* and *Ixodes ovatus* ticks were collected by the flagging method in central Hokkaido, Japan, and morphologically identified [34]. Ticks were then microinjected with approximately 0.18 µL of rYEZV WT or rYEZV GPCΔ28–68 (6.3 FFU/tick) through the anal aperture using an IM-400 microinjector (Narishige). The inoculated ticks were incubated at 16°C for seven days and homogenized in DMEM supplemented with 1% penicillin-streptomycin and 1% L-glutamine. Virus titers were expressed as FFU.

### In silico analysis of YEZV GPC sequence

The full-length amino acid sequence of the YEZV glycoprotein precursor (GPC) was analyzed in silico. The signal peptide was predicted using SignalP 6.0 [35]. Putative transmembrane regions were predicted using SOSUI [36] and TMHMM v2.0 [37]. Potential mucin-type *O*-glycosylation sites were predicted using NetOGlyc 4.0 [38]. All analyses were performed using the respective web servers with default settings.

## Statistical analysis

Statistical analyses were performed using GraphPad Prism 10.6.1 (GraphPad Software). Survival curves were compared using the log-rank (Mantel–Cox) test. Virus titers of tick homogenates were compared using the Mann–Whitney U test.

## Acknowledgements

We thank Ms. Yukari Horio from the Faculty of Veterinary Medicine, Hokkaido University, for technical assistance with tick cell experiments.

## Supporting information

S1 Fig. Signal peptide prediction for YEZV GPC by SignalP 6.0.

S2 Fig. Transmembrane region prediction for YEZV GPC by SOSUI.

S3 Fig. Transmembrane region prediction for YEZV GPC by TMHMM 2.0. S1 Table. *O*-linked glycopeptides identified by Byonic analysis.

S2 Table. Prediction of mucin-type *O*-glycosylation sites in the YEZV GPC. S3 Table. *N*-linked glycopeptides identified by Byonic analysis.

S4 Table. *N*-glycosylation sites in the YEZV GPC identified by IGOT analysis using fully tryptic peptides.

S5 Table. *N*-glycosylation sites in the YEZV GPC identified by IGOT analysis using semi-tryptic peptides.

S6 Table. Unique peptides identified by Mascot analysis. S7 Table. Primers used in this study.

S8 Table. Sequence accession numbers used for phylogenetic analysis.

